# Myasthenia thymus reprograms class-switched B cells into BAFF-dependent survivors

**DOI:** 10.1101/2025.09.19.677075

**Authors:** Yuanqing Yan, Diego Avella Patino, Yimeng Zhao, Xin Wu, G.R. Scott Budinger, Ankit Bharat

**Affiliations:** Department of Surgery, Division of Thoracic Surgery, Feinberg School of Medicine, Northwestern University, Chicago, IL, 60611; DeWitt Daughtry Family Department of Surgery, Miller School of Medicine, University of Miami, Coral Gables, FL, 33124; Department of Medicine, Division of Pulmonary and Critical Care Medicine, Feinberg School of Medicine, Northwestern University, Chicago, IL 60611

## Abstract

Myasthenia gravis (MG) presents a clinical challenge where autoantibody titers against the neuromuscular junction fail to predict disease severity or treatment response, and thymectomy provides inconsistent benefit despite removing the source of autoreactive B cells. We hypothesized that thymic B cells acquire survival mechanisms that bypass normal tolerance checkpoints, enabling persistence independent of antigen-specific selection. Using single-cell RNA sequencing, V(D)J repertoire analysis, and spatial transcriptomics from human thymic samples, representing the largest MG atlas to date (237,661 cells), we discovered a previously unrecognized tolerance checkpoint swap in MG pathogenesis. Pathological class-switched B cells within thymic germinal centers exhibited reduced antigen presentation capacity (decreased CD74, CD1C expression) and diminished CD40 co-stimulation while upregulating TNFRSF17 to engage BAFF survival signals. This shift from T cell-dependent activation to BAFF-driven survival allowed polyclonal autoreactive B cells to persist without stringent selection and seed peripheral sites. T follicular helper cells supported this reprogramming through increased TNFSF13B and CHGB expression, the latter enabling dopamine-mediated acceleration of B cell synapses. These findings resolve longstanding clinical paradoxes in MG and identify the BAFF-BCMA axis as an actionable therapeutic target. Measuring serum BAFF and soluble BCMA could provide superior biomarkers to antibody titers, while BAFF inhibition or CD40 agonism could restore normal tolerance checkpoints, offering new therapeutic strategies for MG and potentially other autoimmune diseases.

**One Sentence Summary:** Myasthenia gravis is characterized by a tolerance checkpoint swap where pathological B cells shift from requiring T cell help and antigen-specific selection to depending on BAFF for survival.

## Introduction

Myasthenia gravis (MG) represents a clinical paradox in which pathogenic autoantibodies against the neuromuscular junction drive muscle weakness and fatigability, yet antibody titers correlate poorly with disease severity and treatment response(*1–4*). Even more puzzling, thymectomy provides inconsistent clinical benefit despite being a mainstay of treatment. This surgical removal the thymus where autoreactive B cells are known to originate should eliminate the source of pathogenic antibodies, yet many patients continued to produce these autoantibodies. These clinical observations challenge our understanding of how autoimmune conditions such as MG are sustained and underscore the inefficacy of current therapeutic approaches, which focus primarily on antibody reduction.

Thymus plays a central yet incompletely understood role in MG pathogenesis. Beyond its classical function in T cell education, the MG thymus can reveal striking pathological changes. Thymic hyperplasia or thymoma occurs in up to 85% of patients(*5*). Ectopic germinal centers form within the medulla, and specialized thymic epithelial cells aberrantly express neuromuscular antigens(*3, 4*). These features point to the thymus not merely as a site of initial tolerance breakdown, but as an active driver maintaining autoimmunity through mechanisms that extend beyond antigen presentation. The persistence of disease despite thymectomy and the disconnect between antibody levels and clinical severity suggest that thymic B cells may acquire survival advantages that allow them to persist independent of conventional selection pressures.

Recent work has identified neuromuscular thymic epithelial cells (nmTECs) that express acetylcholine receptor components and highlighted innate immune signatures in MG thymus(*3–5*). However, these findings alone cannot explain why some B cells escape normal tolerance checkpoints while others do not. They also cannot explain how autoreactive B cells maintain themselves despite therapeutic interventions. Understanding the molecular programs that enable B cell survival within thymic microenvironments could reveal why conventional treatments fail and identify new therapeutic vulnerabilities.

Here, we used a human tissue-anchored integrated multi-omic approach to uncover a previously unrecognized mechanism of tolerance breakdown. We combined single-cell RNA sequencing, V(D)J repertoire analysis, and spatial transcriptomics from the largest MG thymic cohort assembled to date. We demonstrate that MG pathogenesis involves a fundamental reprogramming of B cell survival. Specifically, we identified a shift from classical T cell dependent activation requiring CD40 co-stimulation to an innate-like program driven by BAFF-BCMA signaling. This tolerance checkpoint swap occurs within spatially defined germinal centers and enables diverse autoreactive B cells to survive without stringent antigen specific selection. Our findings resolve longstanding clinical dilemmas in MG and identify actionable therapeutic targets for restoring immune tolerance.

## Results

### Patient characteristics

We collected a total of 23 samples from 16 patients for this study (Table S1). The average age of this cohort is 48.1 years, ranging from 21 to 78 years. The male to female ratio is 5:11. The number of samples for Thymus without MG, Thymus with MG, Thymoma without MG, Thymoma with MG is 3, 8, 3, and 3 respectively. Some patients contributed more than one sample. For example, patient PT4 provided samples of both Thymus without MG and Thymoma without MG. To enhance our analytical power, we also included scRNA-seq data from two external studies: thymoma with MG samples from Yasumizu et al. (includes cells from both peripheral blood and thymoma tissues) and thymic epithelial tumor samples from Xin et al. (*4, 6*). In total, this study included 346,987 cells, of which 237,661 were from our own cohort.

### Single cell data integration and AChR gene expression in thymic epithelial cells

We initiated our analysis by conducting data integration using scvi-tools (Fig. S1A)(*7*), and identified the cell clusters using the Leiden clustering approach. To evaluate the quality of the integrated dataset, we assessed several metrics. Using the scIB package(*8*), we examined batch effect removal and obtained a silhouette score of 0.62, indicating a meaningful reduction in batch effects and reflecting a solid step toward effective data integration. Furthermore, the normalized mutual information (NMI) value of 0.70 demonstrated that the integration effectively preserved the underlying biological structure of the data. We also examined cell distribution bias across datasets within clusters, variations in gene and UMI counts among clusters, and the segregation of cells into distinct populations (Fig. S1B-C). These analyses confirmed that the integration process did not introduce significant biases associated with dataset differences.

Based on the cell compartment marker genes, we successfully identified the major cellular compartment, including epithelial (EPCAM+), endothelial (PECAM1+), immune (PTPRC+) and stromal cells (Fig. S1C). Focusing on structural cells, we leveraged the cell type-specific gene markers to identify 10 epithelial cell types, 6 endothelial cell types and 3 stromal cell types (Fig. S2A). Notably, among epithelial cells, we were able to detect rare populations, such as myoid, ciliated and ionocyte cells from our analysis.

Using the annotated epithelial cells, we investigated the presence of nmTECs, a pathological cell type critical for the ectopic expression of neuromuscular molecules in thymoma with MG, as described by Yasumizu et al.(*4*) We matched cell IDs annotated for this cell type and found that nmTECs did not form a unique cluster but were instead embedded within the KRT15+ mTEC cluster (Fig. S2B). Additionally, we analyzed the expression of genes encoding AChR and found no significant differences between MG and no-MG samples (Fig. S2C). This observation suggests that mechanisms beyond self-antigens may contribute substantially to the pathogenesis.

### Altered B cell immune response in MG patients revealed by pseudo-bulk RNA-seq analysis

Since MG is a B cell–mediated autoimmune disease, we started our analysis from the transcriptomic profile changes in B cells. Using established immune cell gene markers, we identified the B lineage within the immune compartment: B cells, plasma cells, and proliferating B cells (Prolif.B) (Fig. 1A). To compare MG and NoMG groups, we applied a pseudo-bulk analysis with offsets, which offers a scalable and computationally efficient alternative to generalized linear mixed models for controlling the false discovery rate, while maintaining comparable statistical performance(*9, 10*). Our pseudo-bulk RNA-seq analysis identified six significantly downregulated genes in MG patients (Fig. 1B). Notably, *CD1C*, a gene responsible for antigen presentation and interaction with other immune cells (e.g., T cells), was significantly downregulated (Fig. 1C). This suggests that MG patients exhibited reduced immune regulation due to dampened B cell-mediated antigen presentation. Additionally, we found that *LILRA4*, an inhibitory receptor that dampens immune responses via binding with ligands on other cells(*11*), was also significantly downregulated.

**Fig.1.**
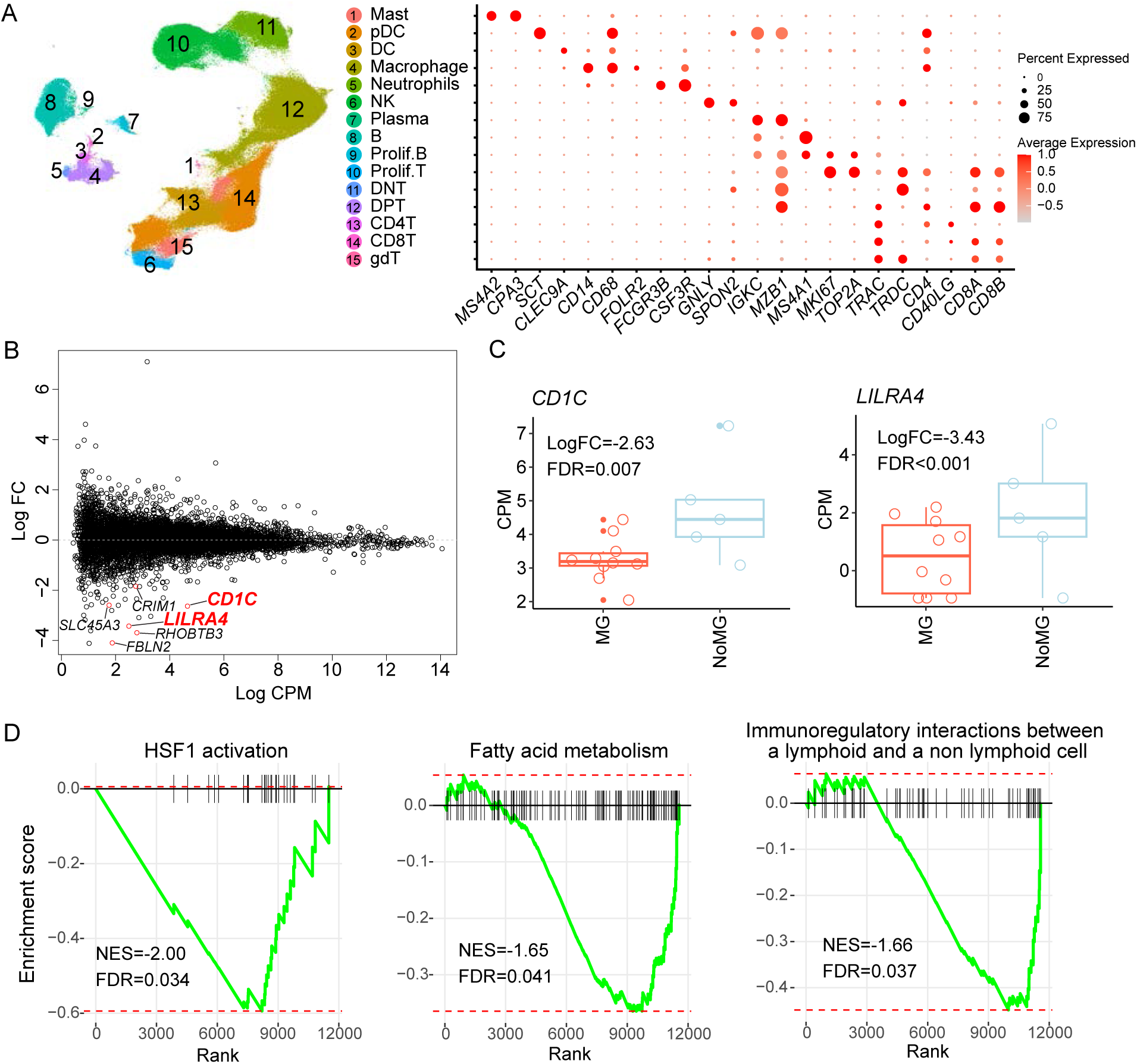
Altered molecular profile in B cells of MG patients revealed by pseudo-bulk RNAseq. (A) UMAP and dot plot showing the identification of B cell lineage from the immune compartment of our integrated object. (B) MA plot delineating the differentially expressed genes in B cells between MG and NoMG. (C) Box plots showing the differential expression of the CD1C and LILR4 genes between MG and NoMG. (D) GSEA plot demonstrating enrichment of three significant pathways in the NoMG samples. Abbreviations: FC: fold change; CPM: Counts per million; NES: normalized enrichment score; FDR: false discovery rate.

Subsequent gene set enrichment analysis (GSEA) revealed multiple biological pathways were significantly dysregulated in MG patients (Fig. 1D). Specifically, the heat shock factor 1 (HSF1) activation pathway was downregulated in the MG group. This pathway is important for the development of regulatory T cell, which help to control inflammation(*12*). We also found a reduction in the fatty acid metabolism pathway. Furthermore, the observed downregulation of immunoregulatory interactions between lymphoid and non-lymphoid cells in the MG group provides additional evidence of disrupted B cell-mediated cellular interactions.

### Identification of pathological class-switching B cells in MG patients

Given the heterogeneity of the B cell lineage, we further examined B cell subtypes using sub-clustering analysis. This revealed three major subtypes—germinal center B cells (GC.B), non-germinal center B cells (no.GC.B), and antibody-secreting cells (ASCs)—defined by the expression of *CCR7, CD38*, and *PRDM1* (Fig. S3A). We then performed fine cell type annotation using cell type–specific markers, identifying a total of 20 distinct B cell subtypes with different enrichment between peripheral blood and tissue, consistent with the findings of Yasumizu et al. (Fig. 2A, Fig. S3B–D). Among these, two subtypes were particularly interesting: the immunoglobulin-class unswitched (labeled as Un_switch) and switched (labeled as Switch) subtypes. The Un_switch resembled a naïve state, with overexpression of the naïve marker *TCL1A*, along with high levels of *IGHD* and *IGHM* (Fig. S3C–D). In contrast, the Switch subtype exhibited high levels of *IGHA1* and overexpression of the memory marker *CD27*. Moreover, both the Un_switch and Switch subtypes are activated B cells, characterized by the overexpression of activation markers *CD83* and *CD69* (Fig. 2B, Fig. S3C). A key molecule that distinguishes these two subtypes is the overexpression of CD70 in the Switch subtype, which fosters B-T cell interactions and drives B cell maturation and differentiation, including plasma cell generation. Notably, both subtypes were detected exclusively in tissue samples and not in peripheral blood (Fig. 2A).

**Fig. 2.**
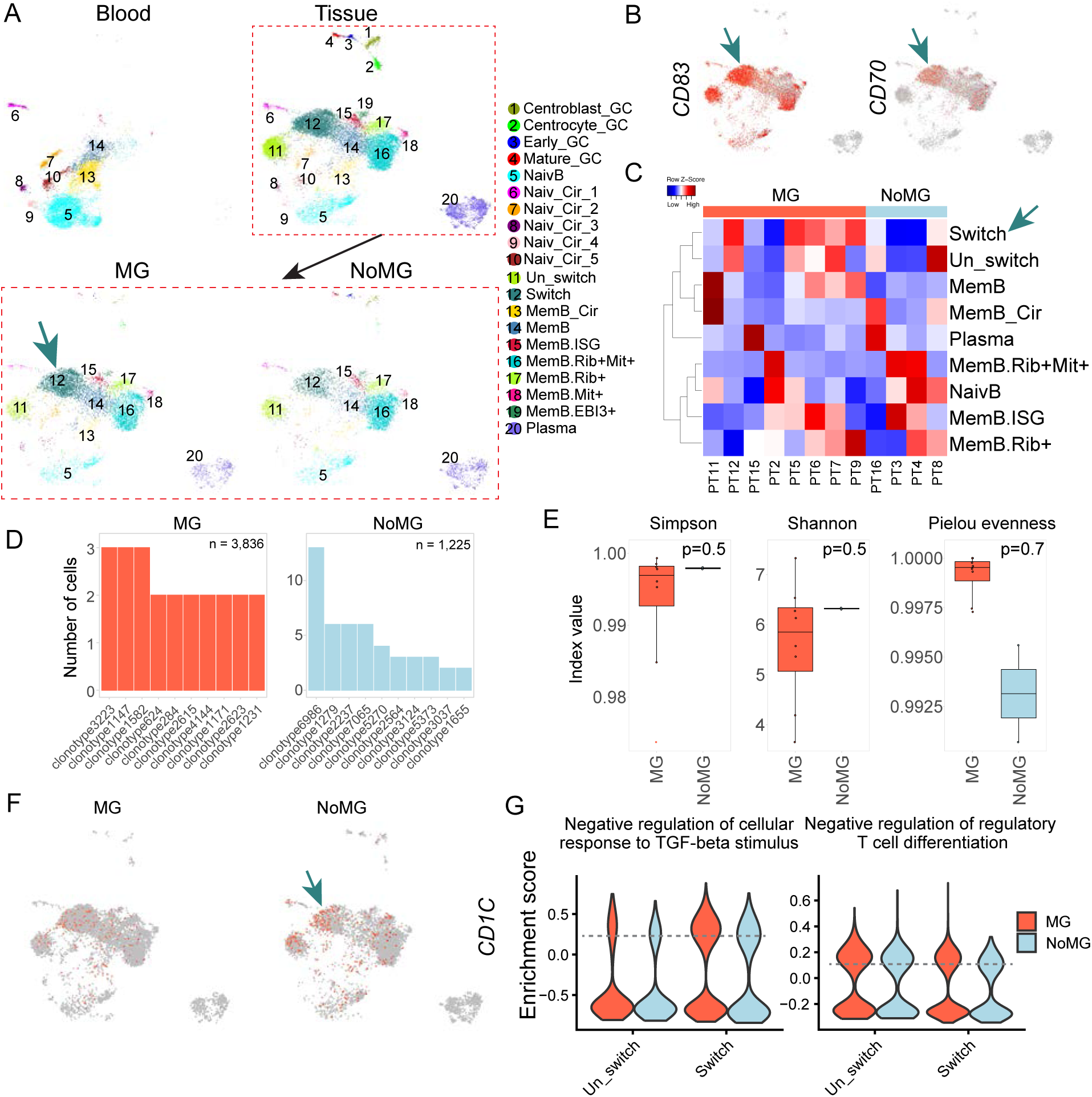
Identification of pathological class-switching B cells in MG patients from single-cell transcriptomic data. (A) UMAP visualization delineating the enrichment of Switch B cells in MG patients. B cell subtypes within the B cell lineage were defined, and UMAPs were split by organ and disease status to illustrate subtype enrichment across conditions. (B) Feature plots showing the overexpression of *CD83* and *CD70* in the Switch B cell subtype. (C) Heatmap demonstrating the proportional distribution of each B cell subtype within the lineage. Proportions were calculated as the number of cells per subtype divided by the total B cell number per patient. (D) Bar plot showing the abundance of the top clonotype within the B cell lineage in MG versus non-MG patients. (E) Box plot showing high B clonotype diversity in MG patients, based on calculated diversity index values. (F) Downregulation of *CD1C* in the Switch B cell subtype, confirming findings from pseudo-bulk RNA-seq. (G) Violin plots illustrating upregulation of two biological process pathways in both Switch and Un-switch B cell subtypes in MG samples.

By splitting our tissue data into MG and NoMG groups, our UMAP revealed a higher abundance of the switched subtype in MG patients compared to the unswitched subtype (Fig. 2A). Cell type proportional analysis, which calculated the proportion of each B cell subtype per sample, confirmed the increased abundance of the switched subtype in MG patients, despite some patient-to-patient variation (Fig. 2C). These results imply that the switched subtype may play a significant role in the microenvironment driving disease progression in MG patients.

Furthermore, our integrated dataset enables tracking of different B cell subtypes across both tissue and peripheral blood. We observed the co-existence of memory B cell subsets, marked by high CD27 expression, in both compartments (Fig. S4). In the NU cohort, RNA velocity analysis indicated that Switch B cells likely serve as precursors to these memory B cells. This suggests that Switch B cells can mature into memory B cells within germinal centers, and that some memory B cells may reside in tissue while also circulating in the blood.

### B cell clonal diversity and dysregulated molecular profiles in MG

To further characterize B cell dynamics, we examined clonal expansion patterns using V(D)J sequencing. Clonotype analysis revealed no dominant B cell clone enriched in MG samples, as indicated by the small number of cells within each top clonotype (Fig. 2D). Repertoire diversity analysis showed comparably high clonotype diversity in the B cell lineage of both MG and NoMG patients, without evidence of clonal expansion (Fig. 2E). This suggests that MG pathology is not driven by a single dominant clone but rather by dysregulated interactions within a diverse B cell pool.

We next investigated molecular differences between MG and NoMG samples. We first confirmed the downregulation of *CD1C*, an antigen-presentation gene identified in our pseudo-bulk RNA-seq analysis, with notably lower expression in the switched subtype of MG patients (Fig. 2F, Fig. S3F). Using GO:BP as the reference, we performed pathway enrichment analysis and identified pathways significantly dysregulated between MG and NoMG within the switched subtype. Notably, the negative regulation of cellular response to TGF-beta stimulus pathway was significantly activated in MG (Fig. 2G), suggesting a loss of tight regulation over the TGF-beta pathway, which is critical for suppressing autoimmunity. The negative regulation of regulatory T cell differentiation pathway was also activated in MG patients. Together, these findings indicate that dysregulated T–B cell interactions, particularly involving the switched subtype, may contribute to the breakdown of immune tolerance and promote autoantibody production.

### Altered expression profile of T follicular helper cells in MG patients

Considering that MG is a T cell–dependent disease and our findings on the role of T cells in B cell interactions, we examined the T cell lineage in our integrated single-cell dataset. Using canonical cell type markers, we identified 24 distinct cell types within the non–B lymphocyte lineage (Fig. S5). Among these, we observed large clusters of double-positive T cells (DPT), double-negative T cells (DNT), and proliferative T cells. Within the CD4+ T cell lineage, we identified multiple regulatory subtypes, including T follicular helper cells (Tfh), regulatory T cells (Treg), and type 1 helper CD4+ effector memory T cells (CD4.Tem(Th1)). When comparing MG and NoMG patients, no T cell subtype was absent from either group (Fig. 3A). This finding was consistent with our cell proportion analysis, which examined the relative abundance of each subtype within the total T cell lineage per sample (Fig. 3B).

**Fig. 3.**
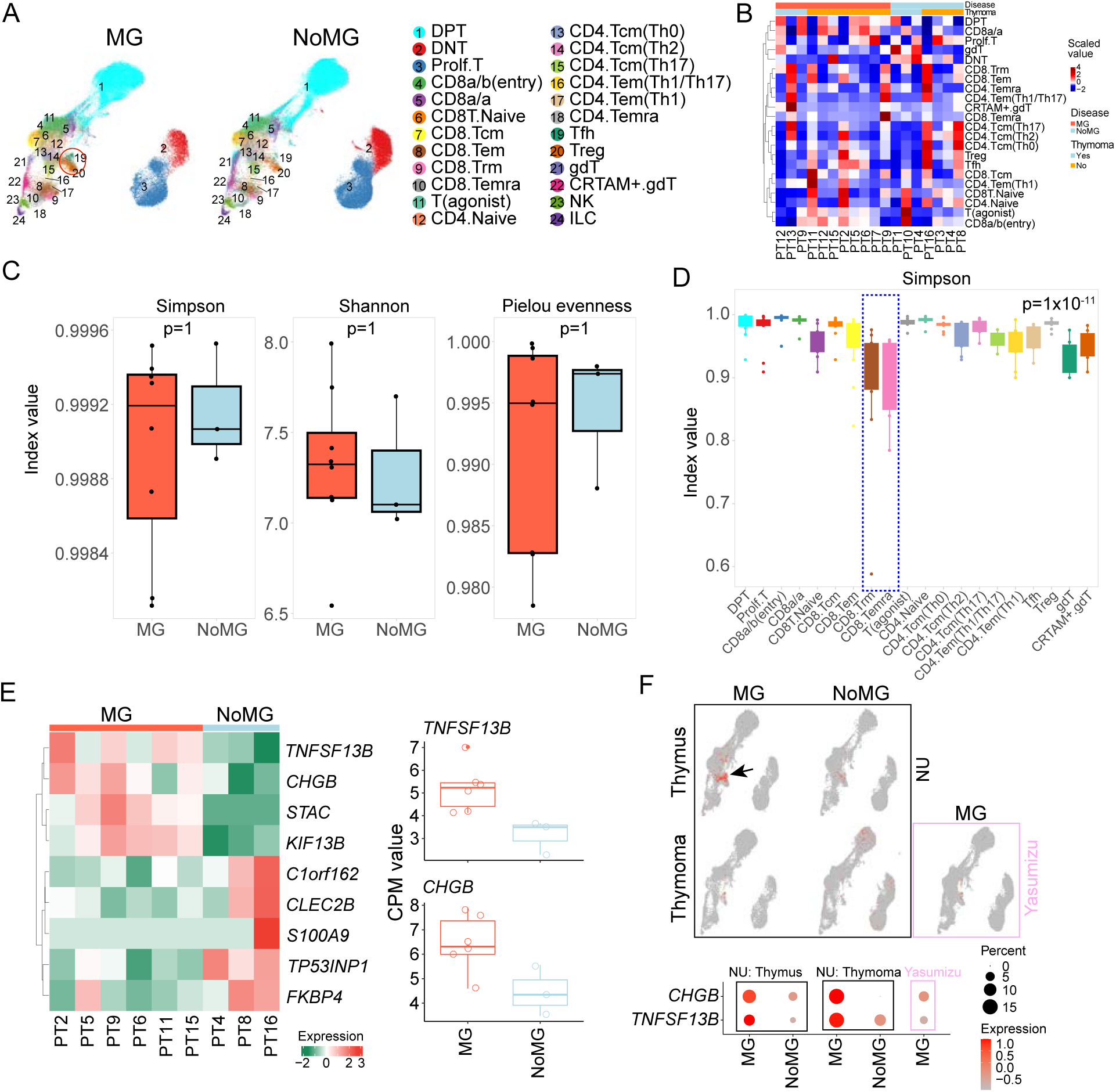
T cell clonal diversity and activation of TNFSF13B and CHGB in Tfh of MG patients. (A) UMAP demonstrating T cell subtype distribution in MG and NoMG samples. (B) Heatmap of proportional distribution of each T cell subtype within the lineage, calculated as the number of cells per subtype divided by total T cells per patient. (C) Box plot showing comparable T clonotype diversity between MG and NoMG patients, based on calculated diversity index values. (D) Relative clonal expansion in CD8+ subtypes (e.g., CD8.Trm and CD8.Temra), indicated by lower Simpson index values. (E) *TNFSF13B* and *CHGB* are significantly upregulated in Tfh of MG patients. Pseudo-bulk RNA-seq was used for differentiation analysis; CPM values were used for heatmap and boxplot plotting. (F) Overexpression of *TNFSF13B* and *CHGB* in Tfh from MG samples was confirmed by scRNA-seq data from both the NU cohort and the Yasumizu study, including thymus with MG and thymoma with MG. Feature plots show *CHGB*’s unique expression in Tfh, rare in other T cell subtypes.

In contrast to the clonotype analysis of B cells, our V(D)J sequencing of the T cell lineage revealed several clonotype expansions. The most expanded clonotype was clonotype337 (109 cells), followed by clonotype1695 (42 cells) and clonotype67104 (29 cells) (Fig. S6A). When stratifying patients into MG and NoMG groups, the overall clonotype fractions were similar, although clonotype337 showed relatively greater expansion in MG patients (Fig. S6B). Furthermore, our repertoire diversity analysis, measuring richness and evenness, revealed no significant differences between MG and NoMG, as supported by various diversity metrics (Fig. 3C). This observation is consistent with the findings from Yasumizu et al.(*4*), indicating no substantial T cell clonotype expansion in MG pathogenesis. Further comparisons across T cell subtypes revealed significantly different between-subtype differences in clonotype expansion (Fig. 3D). Notably, CD8+ T cells, especially CD8.Trm and CD8.Temra, exhibited lower diversity. Analysis of the number of cells per clonotype within each subtype confirmed that expansions were predominantly found in CD8+ T cells, particularly CD8.Trm and CD8.Tem (Fig. S6C). In contrast, CD4+ T cell subsets, including Tfh and Treg, showed no evident clonotype expansions (Fig. S6C).

In light of the significant role of Tfh cells in regulating B cell activity, we focused on Tfh and performed pseudo-bulk RNA-seq analysis comparing MG and NoMG groups. This analysis revealed nine significantly differentially expressed genes, with four upregulated in the MG group (Fig. 3E). Notably, *TNFSF13B*, a molecule crucial for B cell maturation and antibody production, was significantly upregulated in MG patients. Additionally, *CHGB*, which enables Tfh cells to produce and release high amounts of dopamine upon interaction with B cells, was also overexpressed(*13*). Dopamine production has been reported to act as a signaling molecule that strengthens and accelerates T-B interactions, thereby enhancing germinal center output and antibody production—a potential novel mechanism of Tfh involvement in MG pathogenesis(*13*).

In addition to the pseudo-bulk RNA-seq analysis, we validated our findings using scRNA-seq data from both our cohort and public datasets. We confirmed the overexpression of *CHGB* not only in the thymus of MG patients but also in thymoma with MG, as well as in the dataset from Yasumizu et al.(*4*) (Fig. 3F). Notably, *CHGB* overproduction was highly restricted to the Tfh, with little or no expression in other T cell subtypes. Furthermore, our scRNA-seq data also revealed increased *TNFSF13B* expression in Tfh cells, further validating the pseudo-bulk RNA-seq results.

### Altered T–B cell crosstalk revealed by tensor decomposition of cell–cell communication patterns

Although differential gene expression analysis revealed several molecules crucial for MG pathogenesis, it could not fully capture the complex patterns of cell–cell interactions between disease states. Therefore, we performed tensor decomposition of cell–cell communication patterns between T and B cell subtypes to investigate differences in intercellular communication between MG and NoMG samples(*14*). This analysis identified three significant factors (Factors 1, 4, and 8) associated with disease status (Fig. 4A, Fig. S7A). Specifically, examination of signal flow between cell types showed that: Factor 1 represented T–B cell interactions, with signal flow from T cells to B cells; Factor 4 represented T–T cell interactions; Factor 8 represented interactions between B/T cells and T cells, with signal flow from B/T cells to T cells (Fig. 4B). Analysis of Factor 1 revealed that the Switch B cell subtype received strong signals from multiple T cell subtypes, including Tfh, CD4.Tcm(Th17), CD4.Tem(Th1/Th17), and CD4.Tcm(Th2).

**Fig. 4.**
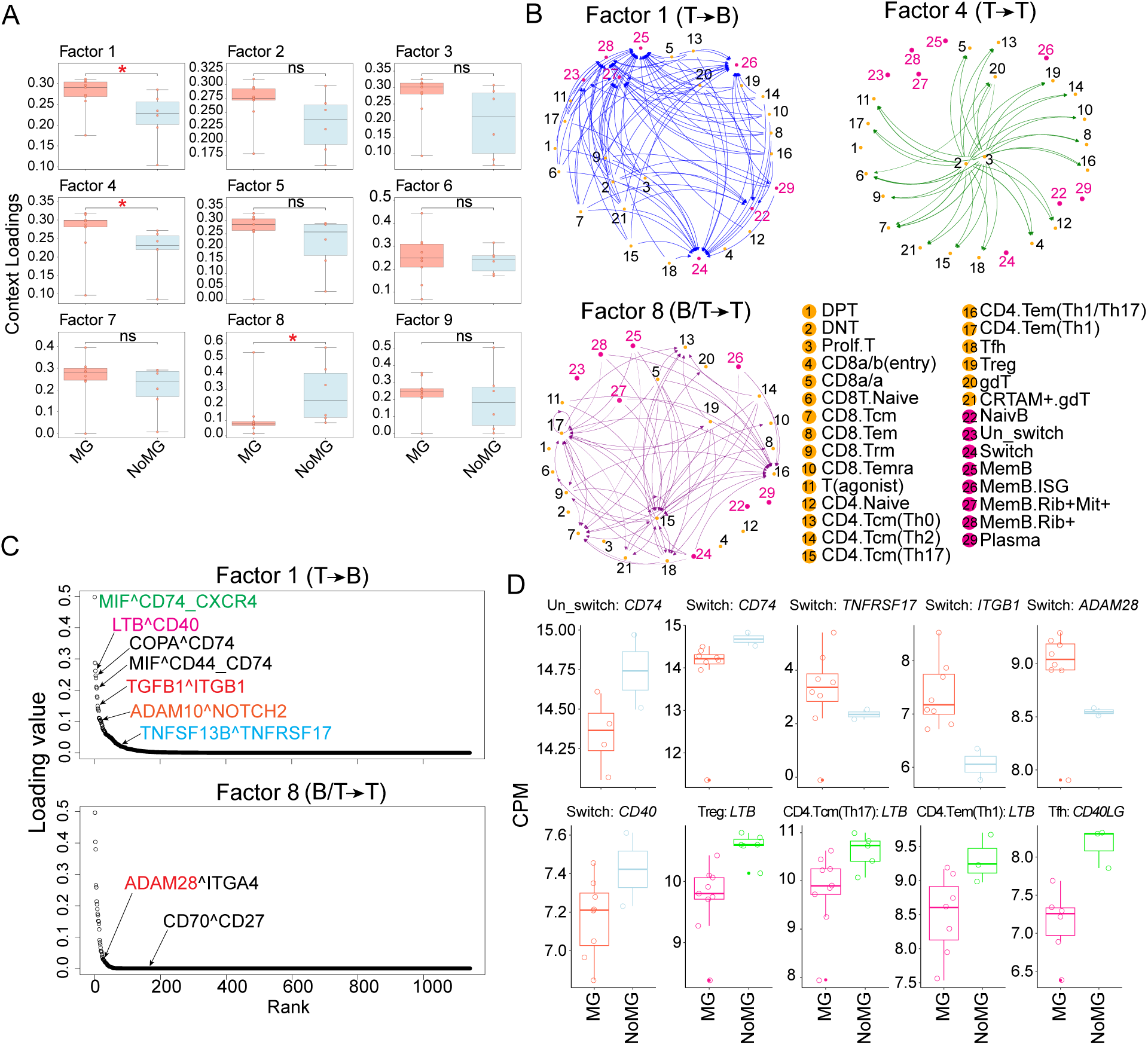
Differential cell–cell communication between MG and NoMG samples. (A) Three factors (1, 4, and 8) were significantly different between MG and NoMG, as shown by context loading values from Tensor-Cell2cell analysis. (B) Network diagrams illustrating signal flow between cell types for the three significant factors. (C) Scatter plot showing the top ligand– receptor pairs in factors 1 and 8. (D) Boxplot of differential expression levels of key ligand/receptor genes between MG and NoMG samples, based on CPM values from pseudo-bulk RNA-seq.

To further explore the specific molecular mechanisms involved, we first analyzed the ligand– receptor pairs in Factor 1. We observed alterations in antigen presentation activities, evidenced by the frequent involvement of *CD74*, a gene crucial for the MHC class II antigen-presentation pathway, among the top-ranked ligand–receptor pairs (Fig. 4C). Analysis of cell types with high *CD74* expression revealed that both Un-switch and Switch B cell subtypes showed the strongest expression (Fig. S7B). However, when comparing MG and NoMG states within these two B cell subtypes, *CD74* expression was markedly reduced in MG patients (Fig. 4D). This finding, together with our previous observation of reduced *CD1C* expression (Fig. 1C, 2F), strongly supports the notion that pathological B cell subtypes in MG have diminished antigen presentation capacity.

Another notable ligand–receptor pair involved the co-stimulatory molecule *CD40* (Fig. 4C), whose binding to its ligand enhances the antigen-presenting capacity of antigen presentation cells. Multiple CD4+ T cell subtypes, including Treg, CD4.Tcm(Th17), and CD4.Tem(Th1), exhibited relatively lower expression of the ligand LTB in MG patients. Correspondingly, Switch B cells in MG patients showed lower expression of the *CD40* receptor. In addition to the LTB–CD40 pair, we also examined the well-known CD40L–CD40 interaction and found the reduced expression of *CD40LG* in Tfh cells (Fig. 4D).

In addition to reduced antigen presentation capacity, we also identified molecular mechanisms that promote B cell activation and adaptation to the microenvironment. One such pathway is the TNFSF13B–TNFRSF17 axis, whose interaction is essential for B cell long-term survival, differentiation, and antibody production. *TNFRSF17* expression was elevated in Switch B cells from MG patients, in parallel with increased *TNFSF13B* production by Tfh cells (Fig. 3E, Fig. 4D). Since *TNFSF13B* is typically produced at high levels by innate immune cells such as monocytes, macrophages, and dendritic cells, we assessed its expression in these myeloid populations and likewise found higher levels in MG samples (Fig. S7C). Together, these findings—reduced LTB/CD40L–CD40 signaling alongside increased BAFF–BCMA (TNFSF13B–TNFRSF17) signaling—suggest that Switch B cells transition from tightly regulated, T-dependent activation mediated by CD40 to an innate-like, BAFF-driven survival program.

Switch B cells not only receive signals from T cells but also produce ligands that feed back to regulate T cell activity. Analysis of Factor 8 revealed that Switch B cells overproduced *ADAM28*, which encodes ADAM metallopeptidase domain 28 and modulates T cell activation via the ADAM28–ITGA4 pathway (Fig. 4C–D). As a molecule critical for promoting lymphocyte adhesion and transendothelial migration, *ADAM28* may contribute to T cell priming and enhanced adhesion, thereby facilitating the delivery of co-stimulatory signals necessary for full T cell activation.

### Spatial transcriptomics identifies Switch B cells within germinal centers

Pathological T–B cell interactions are always associated with the formation of secondary structures(*15*), and understanding such structures in the MG microenvironment—particularly those supporting class-switched B cell activity—could be essential. To investigate this, we performed Visium HD spatial transcriptomics to dissect their spatial distribution and biological activities. Our niche analysis identified two main domains: cortex and medulla/stroma, each characterized by distinct niche-specific gene expression profiles (Fig. 5A). Notably, we observed that B cells were enriched within the medulla/stroma niche.

**Fig. 5.**
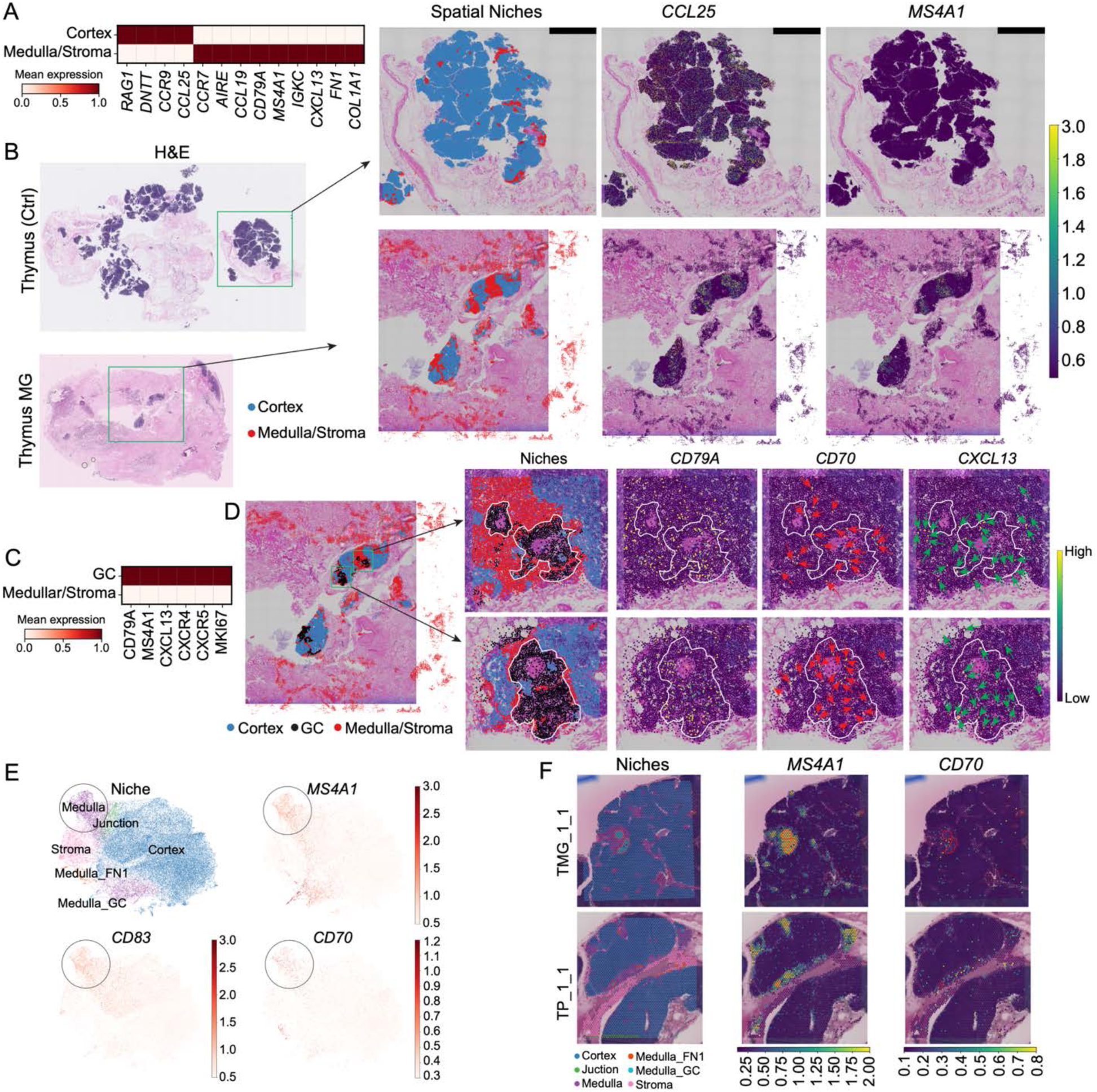
Pathological Switch B cells reside in germinal centers revealed by spatial transcriptomic data. (A) Heatmap showing gene markers defining cortex and medulla/stroma niches from Visium HD spatial transcriptomics of the NU cohort. (B) Spatial distribution of niches in thymus with and without MG from Visium HD spatial transcriptomics. *CCL25* marks the cortex niche, and *MS4A1* marks B cells. (C) Heatmap showing gene markers defining germinal centers within medulla niches. (D) Switch B cells reside within and around germinal centers in MG patients, evidenced by co-localization of *CD70*. (E) UMAP showing residence of Switch B cells in medulla niches from Visium spatial data in the Yasumizu study(*3*). (F) Localization of Switch B cells within and around germinal centers was further confirmed in two MG samples from the Yasumizu study.

Compared with the NoMG sample, the MG sample showed a significant enrichment of B cells forming spatial clusters within the medulla, suggesting the presence of germinal centers (GCs) (Fig. 5B). Our analysis of the medulla/stroma compartment pinpointed GCs, evidenced by the overexpression of GC-specific gene markers (Fig. 5C). Furthermore, the spatial mapping of Switch B cells revealed that they were mainly located within and surrounding these GCs in the MG sample (Fig. 5D), suggesting that GCs are a structural niche for their maturation and functional activity.

In addition to our in-house data, we cross-referenced Visium spatial transcriptomics from Yasumizu et al(*3*). UMAP analysis of this external data also showed Switch B cells within the medulla niche (Fig. 5E). Consistent with our own findings, these cells were found surrounding GCs in MG patients (Fig. 5F).

### Spatial co-localization of key ligand–receptor pairs within and around the GCs

To investigate the aberrant ligand-receptor interactions that lead to the breakdown of immune tolerance in MG, we examined their spatial distribution using our spatial transcriptomic data. First, we investigated the spatial distribution of *CD27*, a receptor that binds to *CD70*. We observed that *CD27* co-localizes with GCs in both our Visium HD and the external spatial data (Fig. 6A-B). Since Switch B cells overexpress receptors like *TNFRSF17* to promote long-term survival by interacting with Tfh cells (Fig. 4C-D), we next examined the spatial expression of *TNFRSF17*. Our findings show that GCs provide the microenvironment necessary for this overexpression, which in turn mediates Switch B cell activity (Fig. 6C-D). Additionally, our spatial transcriptomic data confirmed the co-localization of *ADAM28*—a ligand overexpressed by Switch B cells to interact with T cells—within the GCs (Fig. 6E-F). Together, these results indicate that GCs serve as specialized niches supporting the survival and functional reprogramming of pathological B cells, ultimately driving the breakdown of immune tolerance in MG pathogenesis.

**Fig. 6.**
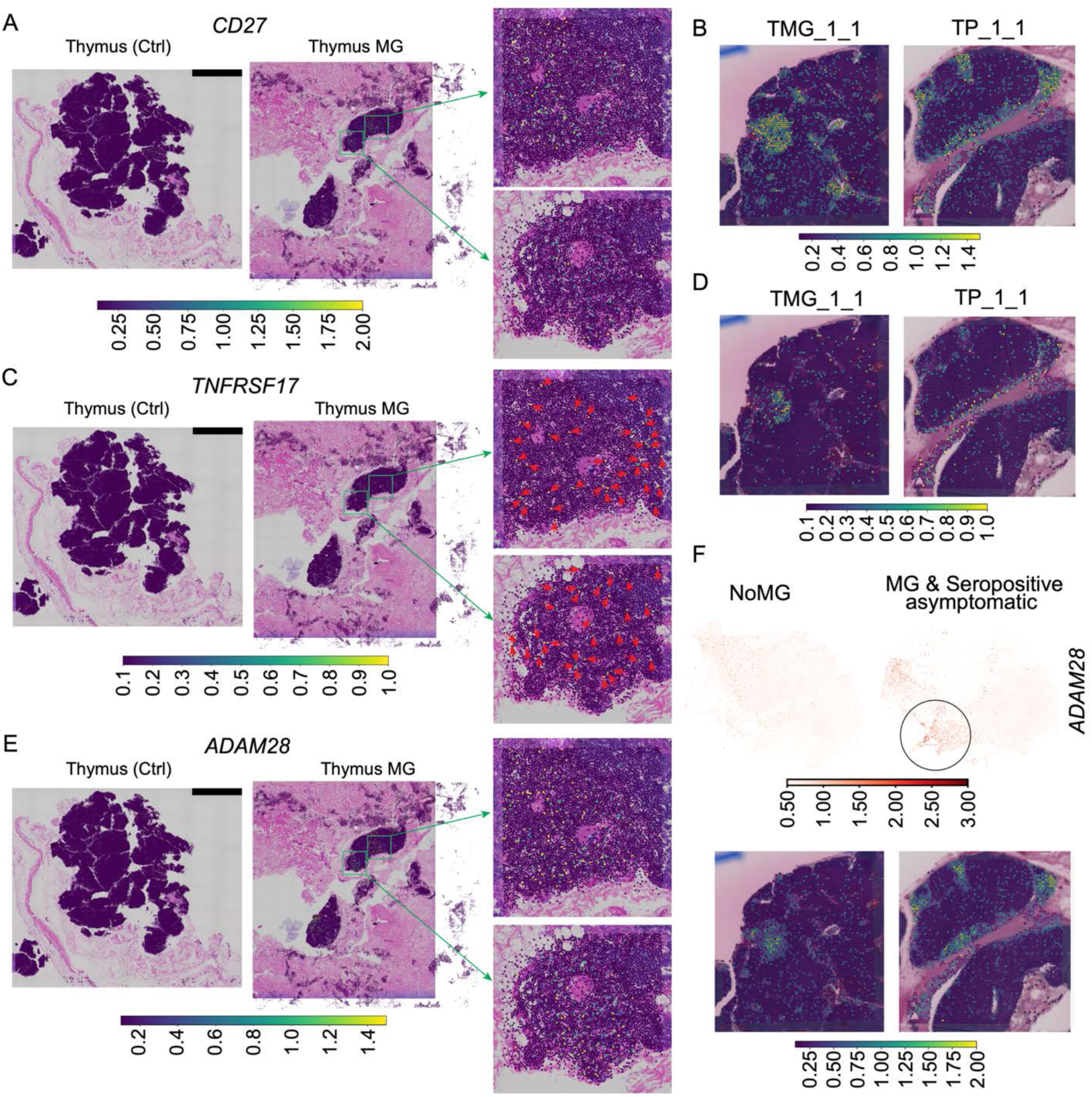
Germinal centers as key structures mediating aberrant cell–cell communication in MG pathogenesis. (A, C, E) Spatial distribution of *CD27* (receptor of *CD70*), *TNFRSF17* (receptor of *TNFSF13B*), and *ADAM28*, respectively, showing their expression within and around germinal centers in MG patients from the NU cohort (Visium HD spatial transcriptomics). (B, D, F) Validation of the spatial distribution of *CD27, TNFRSF17*, and *ADAM28*, respectively, within and around germinal centers using publicly available Visium spatial transcriptomic data.

## Discussion

This study suggests a fundamental shift in myasthenia gravis pathogenesis. Rather than resulting from simple autoantigen exposure or clonal expansion of autoreactive B cells, MG arises from a reprogramming of immune tolerance checkpoints within the thymus. We demonstrate that pathological B cells escape normal selection pressures by switching from classical T cell dependent activation to an innate survival program driven by BAFF signaling(*16–18*). This tolerance checkpoint swap represents a new paradigm for understanding not only MG but potentially other autoimmune diseases where clinical severity disconnects from autoantibody levels(*15, 19*).

Our data suggests that class switched B cells that acquire unique survival advantages within thymic germinal centers represents a key mechanism that drives autoimmunity. These cells exhibit three critical features that distinguish them from normal B cells. First, they show markedly reduced antigen presentation capacity through downregulation of MHC class II genes including CD74 and CD1C. Second, they demonstrate diminished CD40 expression and reduced responsiveness to CD40L from T cells. Third, they compensate for these deficits by upregulating TNFRSF17 (BCMA) to engage BAFF survival signals. This molecular reprogramming allows B cells to survive without stringent antigen specific selection. Indeed, the polyclonal nature of the B cell repertoire we observed supports this model. Rather than clonal expansion of high affinity autoreactive cells, MG involves the survival of diverse B cell populations that would normally be eliminated through tolerance checkpoints.

This mechanism resolves several longstanding clinical observations in MG, for example, the discordance between antibody titers and disease severity. Antibody levels measure only the output of plasma cells, while disease activity potentially reflects the strength of survival signals maintaining the entire autoreactive B cell pool. Patients with high survival tone in their thymic niches may experience severe disease even with modest antibody titers. Conversely, those with lower survival signaling may have high titers but milder symptoms. The variable efficacy of thymectomy also becomes explicable. Class switched B cells that depend on BAFF rather than antigen specific signals can seed peripheral sites before surgery as suggested by our data of tissue and blood B cells. These cells can establish extrathymic reservoirs that persist after thymic removal. Our spatial transcriptomics data showing ADAM28 expression further suggests these cells acquire tissue adhesion programs that may facilitate their peripheral establishment.

The identification of BAFF as a central driver has broader implications for autoimmunity. BAFF overproduction occurs in multiple autoimmune diseases including systemic lupus erythematosus and rheumatoid arthritis(*20*). Our findings suggest that BAFF may enable a common mechanism whereby autoreactive B cells circumvent normal tolerance checkpoints. The parallel upregulation of CHGB in T follicular helper cells provides additional insight. This molecule enables dopamine release at immune synapses, accelerating B cell activation independent of antigen affinity(*13*). Together, BAFF and dopamine signaling create a permissive environment where quantity of survival signals overrides quality of antigen recognition. This framework may explain why BAFF inhibition shows efficacy across diverse autoimmune conditions despite different target antigens. Our findings identify multiple therapeutic opportunities beyond current approaches. First, measuring serum levels of soluble BAFF and BCMA byproducts could provide better biomarkers than antibody titers for monitoring disease activity and predicting treatment response as these markers directly reflect the survival tone driving pathogenic B cells. Second, the BAFF-BCMA axis represents an actionable target. Belimumab, an FDA approved BAFF inhibitor, warrants investigation in MG clinical trials. BCMA targeted therapies under development for multiple myeloma could be repurposed for severe MG cases. Third, restoring CD40 signaling through CD40 agonists might reestablish normal selection checkpoints. This approach could eliminate low affinity autoreactive cells while preserving protective immunity. Fourth, blocking ADAM28 mediated adhesion could prevent B cells from establishing peripheral reservoirs. Finally, combination strategies targeting both survival signals and adhesion programs may prove more effective than single agent approaches.

The spatial organization we identified suggests additional interventions. Germinal centers in MG thymus create specialized niches where pathological signaling concentrates. Disrupting these structures pharmacologically could eliminate the protective microenvironment sustaining autoreactive cells. The colocalization of switch B cells with specific T cell subsets also suggests that targeting their interaction zones could be therapeutic. Agents that modulate germinal center formation or maintenance deserve investigation in MG.

This study has certain limitations. The cross-sectional design prevents us from determining whether the tolerance checkpoint swap initiates disease or develops secondarily. Longitudinal studies examining thymic changes from disease onset through progression will be necessary to establish causality. Our sample size, while the largest single cell atlas of MG to date, limits our ability to detect differences between antibody subtypes. Future studies should specifically recruit patients with MuSK, LRP4, and seronegative MG to determine whether different antibody specificities associate with distinct survival programs. The functional consequences of reduced antigen presentation in Switch B cells also require direct experimental validation through ex vivo coculture systems.

In conclusion, we have identified a tolerance checkpoint swap that provides new insights into MG pathogenesis. The shift from antigen dependent selection to BAFF driven survival explains clinical paradoxes in MG. Importantly, this mechanism identifies specific molecular targets for therapeutic intervention. By restoring the balance between survival signals and selection checkpoints, we may be able to reestablish immune tolerance without broad immunosuppression. Hence, these findings could improve treatment for MG and potentially other autoimmune diseases driven by similar mechanisms.

## Material and Methods

### Human sample preparation and single-cell sequencing

Human tissue samples were obtained from Northwestern Memorial Hospital in Chicago, IL. Two library preparation methods were used: 10x Genomics Chromium Single Cell 3’ v3 chemistry and Single Cell 5’ chemistry with V(D)J sequencing. Four samples were prepared using the 3’ v3 method at the Metabolomics Core Facility at Northwestern University (NU). Tissues were dissociated into single-cell suspensions and sorted via flow cytometry to enrich for epithelial cells (CD45-EPCAM+), immune cells (CD45+), and other cells (CD45-EPCAM-). Each compartment was then subjected to single-cell sequencing. The left nineteen samples were processed with the Chromium Single Cell 5’ chemistry with V(D)J sequencing at the NuSeq Core at NU. Similar dissociation and sorting procedures were applied, but the three compartments were recombined for each sample in varying proportions to prevent overrepresentation of immune cells and ensure balanced representation across cell types. For both methods, sequencing was performed on an Illumina platform, and raw data were demultiplexed with Cell Ranger using standard protocols. Sequencing reads were aligned to the human reference genome (GRCh38) to generate raw gene expression counts per cell.

### Single-cell data quality control and integration

scRNA-seq data were first processed with the SoupX toolkit to remove ambient RNA contamination(*21*), followed by doublet detection and removal using scDblFinder(*22*). Additional quality control was performed with Seurat(*23*); cells were excluded if they had fewer than 200 or more than 7,500 detected genes, fewer than 400 or more than 40,000 UMI counts, or greater than 10% mitochondrial gene content. To increase statistical power, we also incorporated two publicly available datasets: thymoma with MG samples from Yasumizu et al. and thymic epithelial tumor samples from Xin et al.(*4, 6*)

Raw counts were used for data integration with scvi-tools(*7*), a deep probabilistic framework for single-cell omics analysis that leverages GPU acceleration with mini-batching to efficiently model complex datasets while correcting for technical variation. Each sample was treated as a distinct batch, and the top 3,000 highly variable genes were selected to support dimensionality reduction and integration. The integrated latent space was computed using UMAP based on scVI-inferred embeddings and used for downstream analyses. Integration quality was evaluated by examining cell distribution across datasets within clusters, variation in gene and UMI counts among clusters, and separation of distinct cell populations. We further applied the scIB package to compute evaluation metrics(*8*): the silhouette score was used to assess batch correction by measuring how well cells from the same batch clustered relative to cells from different batches, while normalized mutual information (NMI) was used to assess biological conservation by comparing clustering results with annotated cell labels.

### Cell type annotation

We first classified cells into epithelial, endothelial, immune, and stromal compartments using compartment-specific gene markers (*EPCAM*, *PECAM1*, and *PTPRC*), and then extracted cells from each compartment for fine-resolution sub-clustering. Cell type annotation was performed using CellTypist in combination with manual examination(*24*). For the CellTypist analysis, we applied the well-annotated reference labels from a previous study as well as the Immune V2 dataset from the Cell Type Encyclopedia(*4, 25*). For manual annotation, we conducted differential gene expression analysis between clusters to identify cluster-specific markers, iteratively adjusting the clustering resolution to best capture cell type distinctions and align with known populations. During annotation, clusters containing doublets or low-quality cells were removed. Where possible, our annotations were further cross-validated against the original published datasets to ensure consistency and robustness.

### Pseudo-bulk RNAseq analysis

For the pseudo-bulk RNA-seq analysis, we extracted the relevant cell types from the integrated object and summed the raw counts of all cells per sample to enable patient-level analysis. Only samples with cell numbers above the threshold (e.g., n = 25 for B-cell pseudo-bulk RNA-seq) were retained. Differential gene expression analysis was performed using the edgeR package(*26*), restricting the analysis to protein-coding genes. A linear model accounting for disease status and sequencing chemistry was fitted, followed by post-hoc comparisons to identify differential expression between MG and NoMG. False discovery rates (FDR) were estimated using the Benjamini–Hochberg procedure. Genes with FDR < 0.05 and an absolute log2 fold change > 1 were considered statistically significant. GSEA was performed using the GO:BP database from MSigDB as the reference(*27*), implemented with the fgsea R package. Pathways with adjusted p values < 0.05 were considered statistically significant.

### Cell subtypes proportion analysis

We calculated the proportion of each cell subtype within its parent cell lineage for every sample. For example, within the B-cell lineage, the relative abundance of each B-cell subtype per sample was used for hierarchical clustering analysis. Patients with insufficient cell numbers and cell types with low total counts were excluded (e.g., patients with <50 cells and B-cell subtypes with <200 total cells). To ensure consistency, the proportion analysis was performed using tissue data from the NU cohort.

### Bioinformatics analysis of V(D)J sequencing data

For the BCR/TCR sequencing data, we used the djvdj package, including only cells with high-quality transcriptomic data. Samples with fewer than 10 cells were excluded to ensure robustness. Clonotypes were defined based on CDR3 amino acid sequences, and only clonotypes with exactly two chains were retained for further analysis. Cells without an assigned clonotype were removed. The final dataset for the B-cell lineage included 5,061 cells (1,225 from NoMG patients and 3,836 from MG patients). For the TCR data, 19,602 cells were analyzed (4,917 from NoMG and 14,685 from MG patients). Repertoire diversity was estimated using bootstrapping with 500 repetitions and a down-sampling approach to calculate diversity indices.

### Differential cell-cell communication between MG and NoMG samples

Differential cell–cell communication analysis was performed using the LIANA+ package(*28*), with the Tensor-Cell2cell analytical framework applied for intercellular context factorization(*14*). Specifically, we extracted B- and T-cell subtypes from the tissue data of the NU cohort in our integrated object and excluded cell types with fewer than 200 cells from further analysis. We first inferred consensus-based rankings of cell–cell interactions for each individual sample, followed by tensor decomposition to investigate the factors and their associated loadings. The weights of ligand–receptor interactions contributing to each factor were then ranked for subsequent analysis. Statistical significance was defined as an adjusted *p*-value < 0.05.

### Visium HD spatial transcriptomics data analysis

We analyzed clinical and single-cell transcriptomic data to select three representative patients-from a thymus without MG, a thymus with MG, and a thymoma with MG (data not shown)-for Visium HD spatial transcriptomic analysis. Formalin-fixed, paraffin-embedded (FFPE) specimens were sectioned at 5 μm and processed at the NUSeq Core following the manufacturer’s protocol. Images were manually aligned using Loupe Browser (v8.0.0), and genome mapping was performed with Space Ranger (10x Genomics) against the human reference genome (GRCh38). To reconstruct cell-level resolution, the Bin2cell package was used to aggregate 2 μm bins based on H&E morphology and gene expression, at a resolution of 0.5 μm per pixel(*29*). Downstream analyses were conducted with Scanpy(*30*), excluding cells with fewer than 100 or more than 40,000 features, or with mitochondrial content greater than 20%. Cell types were inferred using CellTypist with a confidence score threshold of 0.7, by referencing our scRNA-seq dataset to ensure consistent annotations(*24*).

Spatial niches were initially identified based on predicted cell types and their corresponding spatial coordinates. These niches were further annotated according to their gene expression patterns, such as the overexpression of *CCL25* in the cortex and *CCL19* in the medulla. To identify germinal centers, we focused on the medulla/stromal niche and applied CellCharter to delineate the spatial domains characteristic of these structures(*31*).

## Acknowledgement

We sincerely thank the patients who participated in this study. The scRNA-seq data were processed at the NUSeq Core and the Metabolomics Core Facility at NU, and we are grateful for their assistance. We also thank the NUSeq Core for their support in processing the Visium HD spatial transcriptomics sequencing.

## Supplementary Information

**Figure S1.**
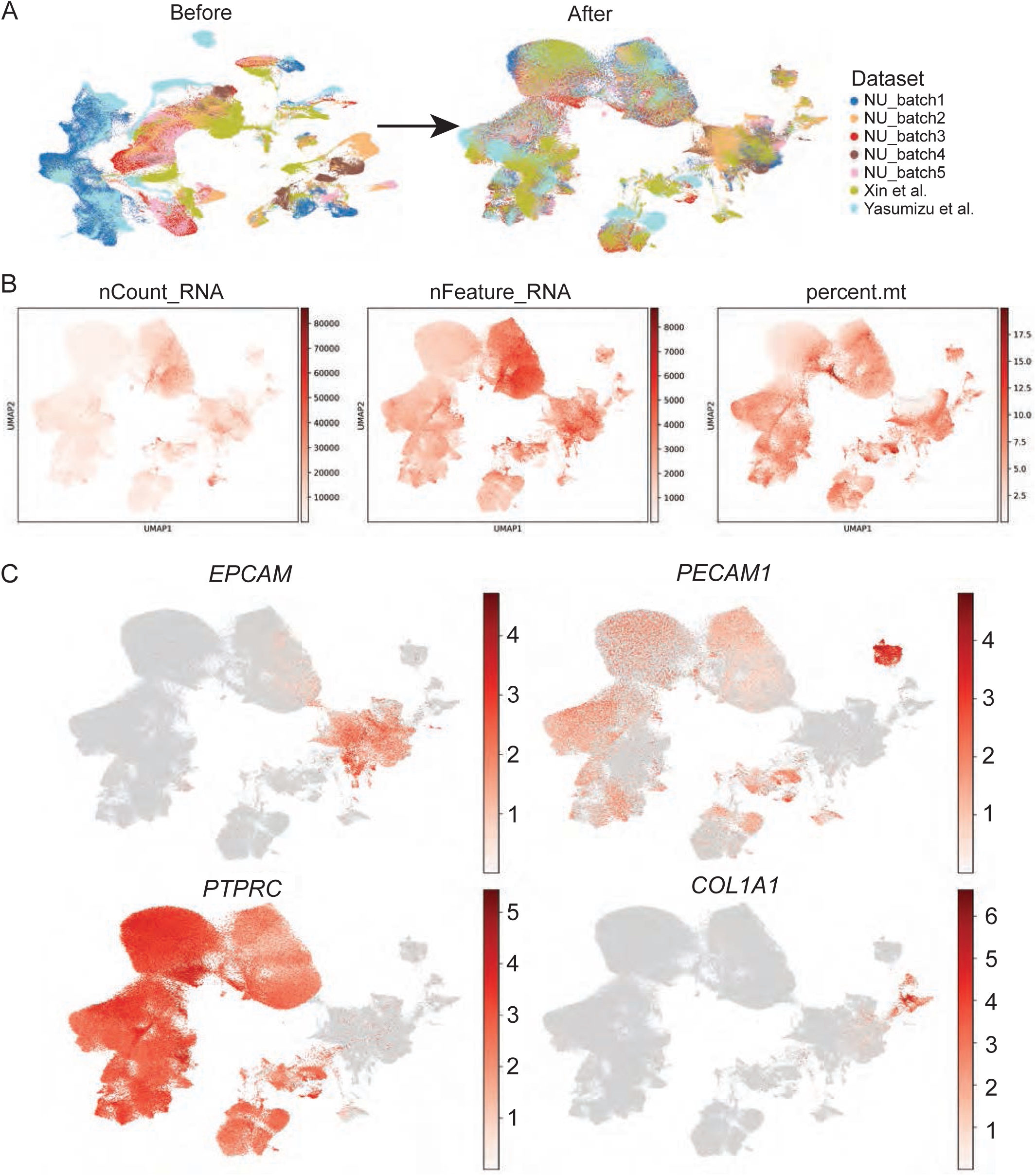
Quality control of single-cell transcriptomic data integration. (A) UMAP plots illustrate cell clustering before and after data integration using scvi-tools. The successful harmonization of cells from different batches indicates high-quality data integration. (B) Distribution of gene count, UMI count and mitochondrial gene percentages across cell clusters. (C) Four cellular compartments were identified based on the expression levels of *EPCAM*, *PECAM1*, and *PTPRC*.

**Figure S2.**
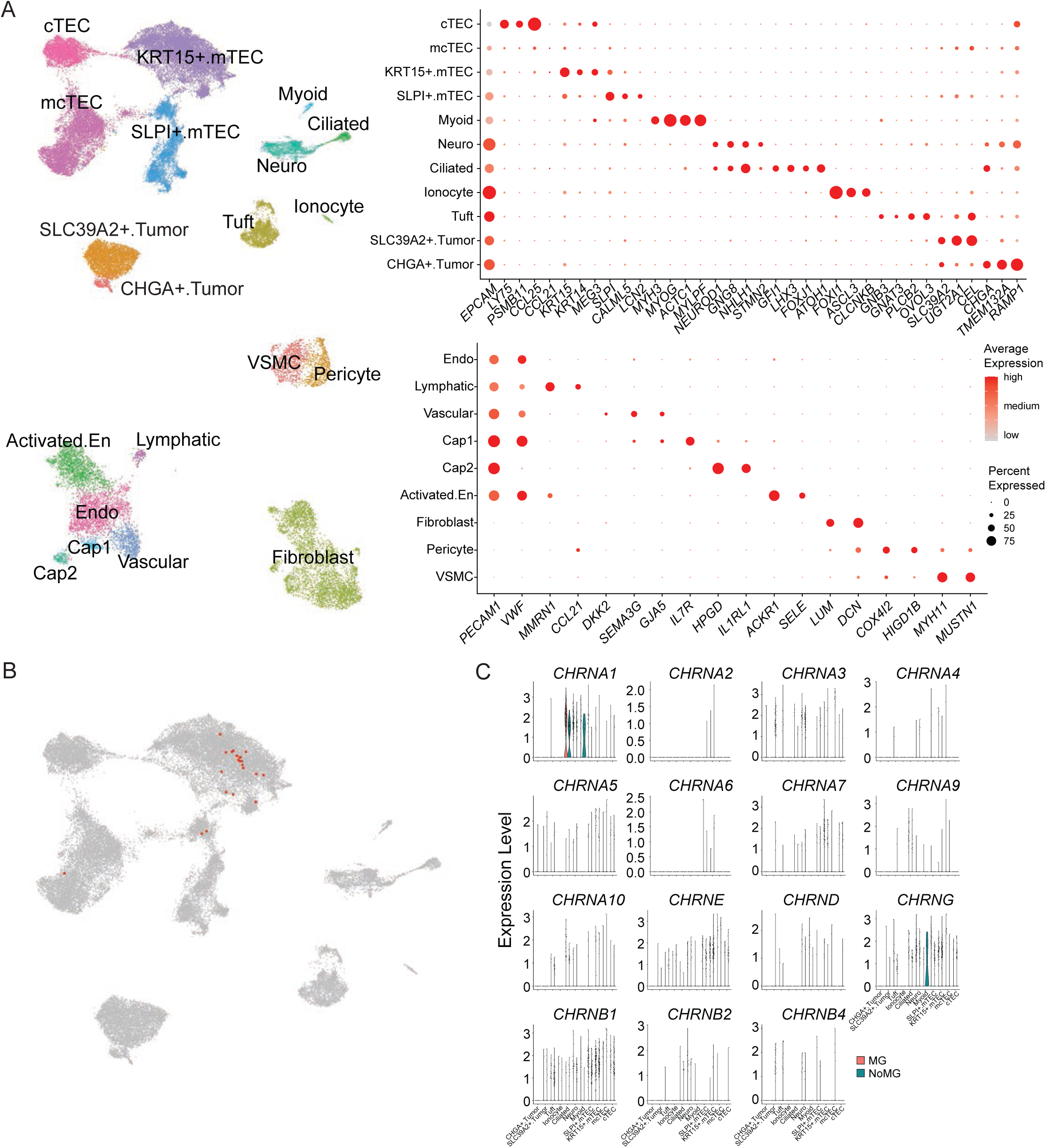
Identification of structural cell populations and AChR gene expression in epithelial cells between MG and NoMG. (A) UMAP visualization of structural cell populations with cell type–specific markers illustrated by a dot plot. (B) Feature plot showing the presence of nmTECs, matched to cell IDs reported in the Yasumizu study. (C) Violin plot showing the expression of AChR genes in epithelial cells from MG and NoMG samples.

**Figure S3.**
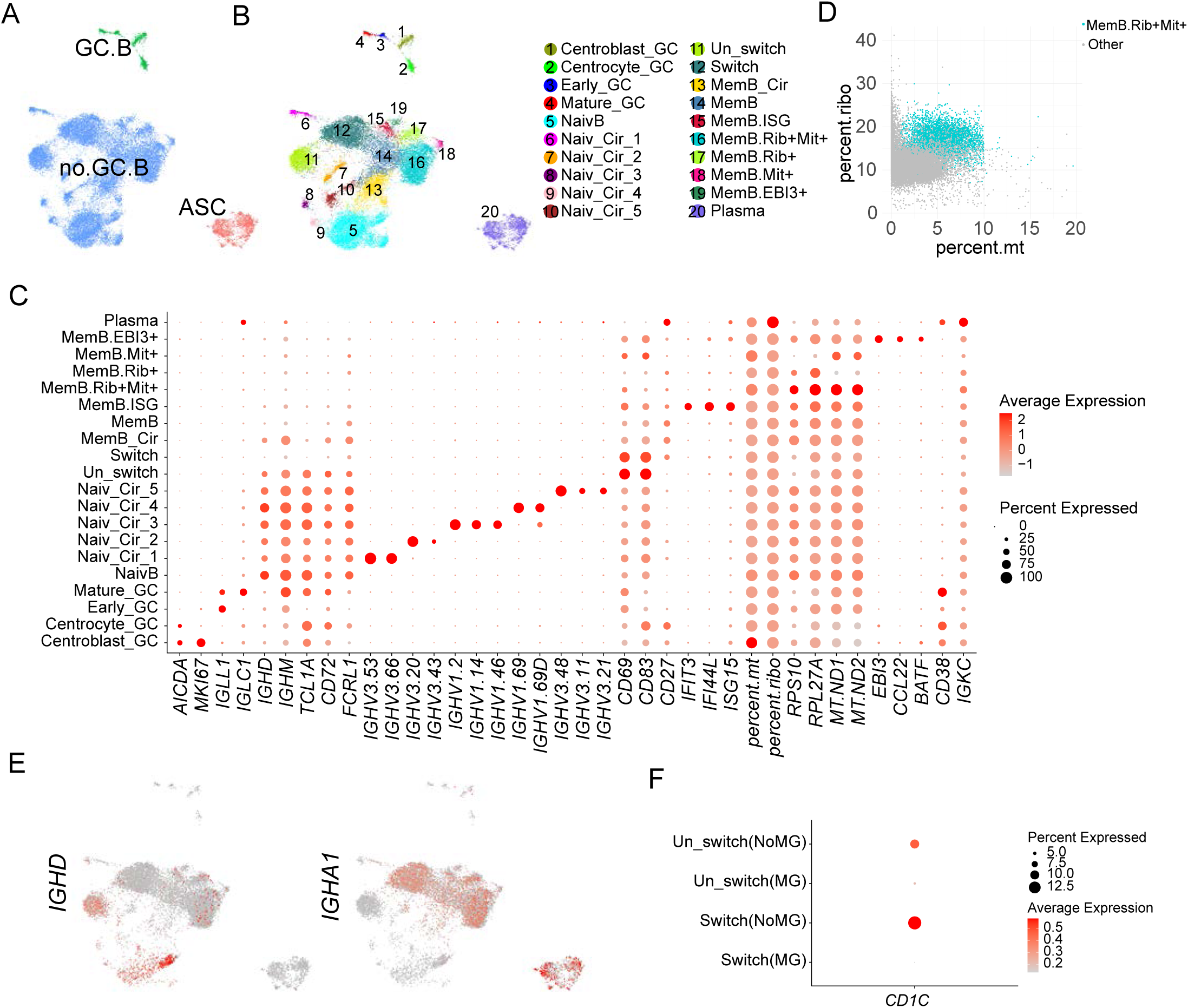
Identification of B cell subtypes within the B cell lineage. (A) UMAP visualization of B cell subtypes identified in this study, including three major subtypes and twenty fine subtypes. (B) Dot plot showing gene markers for each B cell subtype. (C) Scatter plot showing that some B cell subtypes can be defined by mitochondrial and ribosomal gene content. (D) Feature plot showing differential expression of IGHD and IGHA1 between Switch and Un-switched B cells. (E) Dot plot showing reduced expression of CD1C in MG samples.

**Fig. S4.**
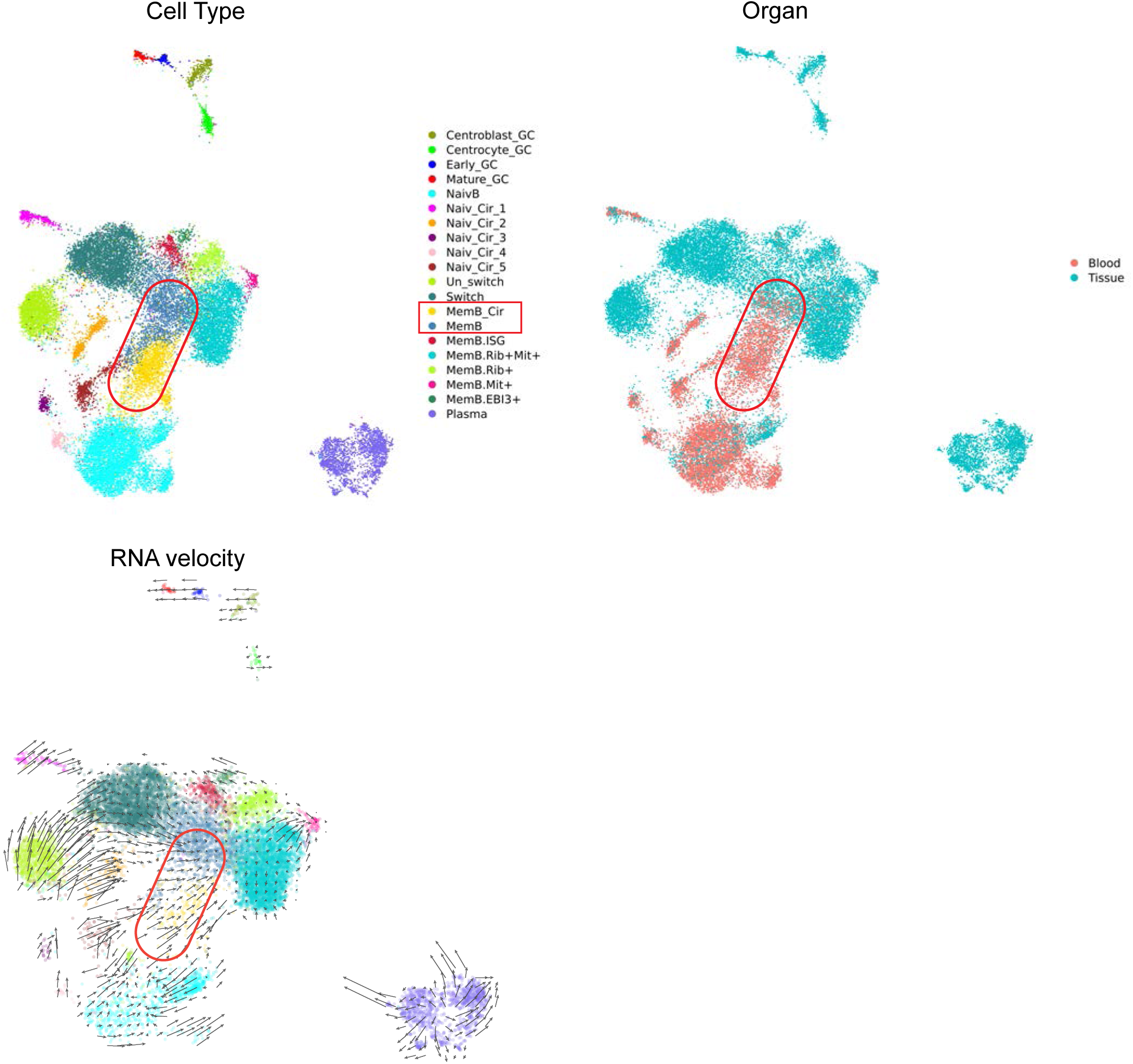
Coexistence of circulating and tissue-resident memory B cells in MG. Circulating and tissue-resident memory B cells are highlighted on the UMAP, and RNA velocity analysis of the NU cohort was performed to illustrate the differentiation of Switch B subtype into memory B cells.

**Figure S5.**
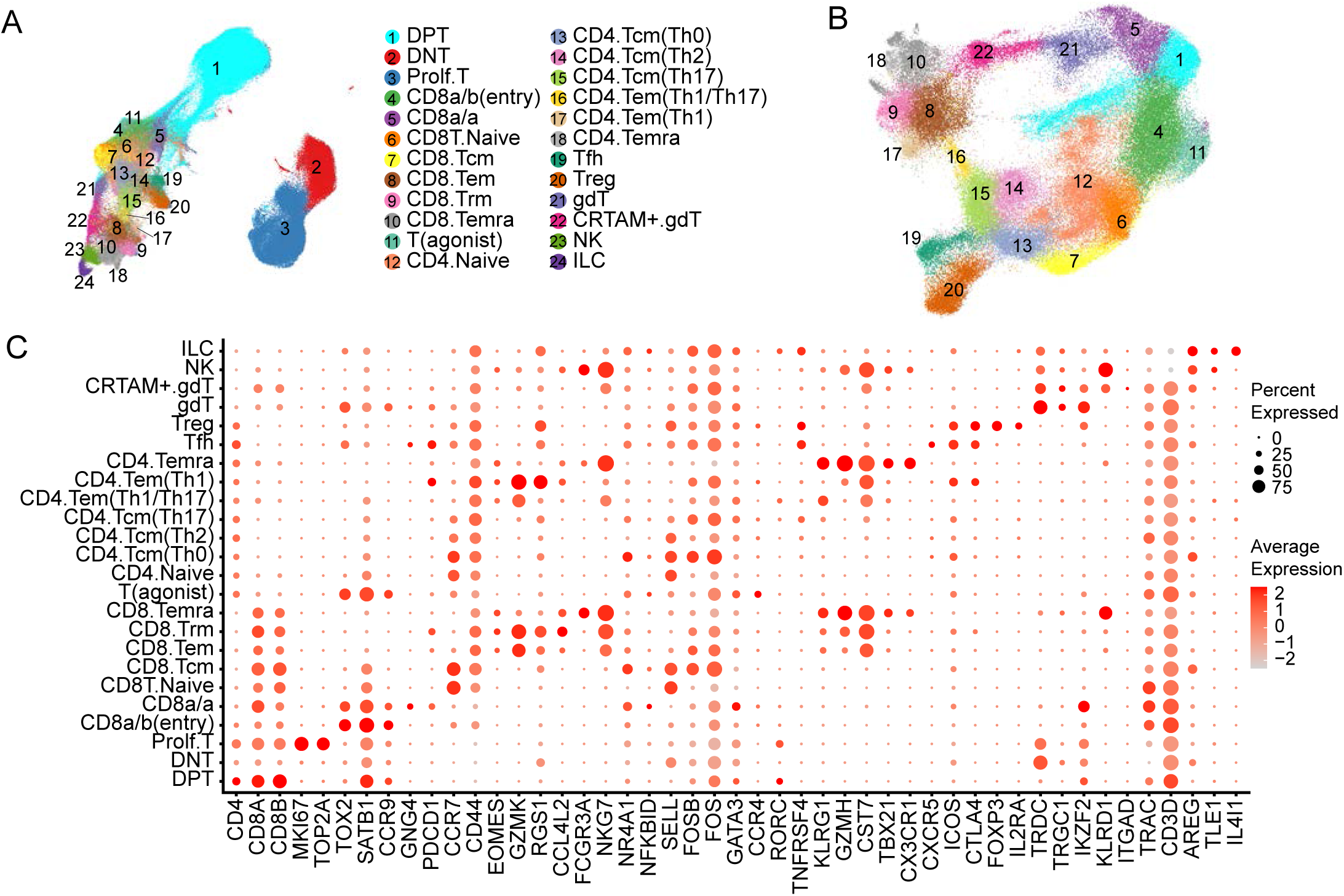
Identification of T cell subtypes within the T cell lineage. (A) UMAP visualization of 24 T cell subtypes identified in this study. (B) UMAP visualization after reclustering, with highly abundant DNT and Proliferating T cells, most DPT cells, as well as NK and ILC cells removed. (C) Dot plot showing gene markers for each T cell subtype.

**Figure S6.**
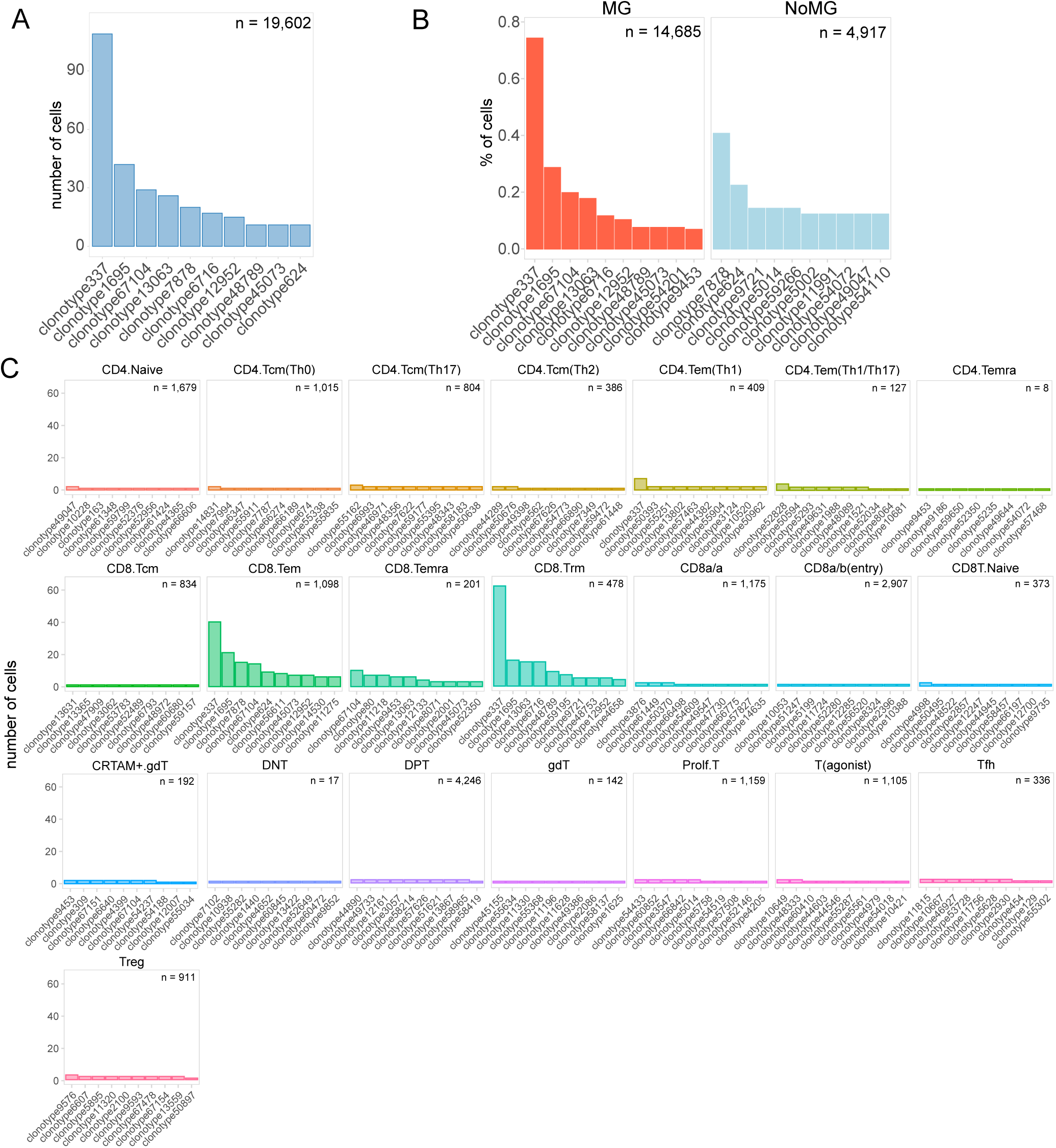
T cell clonal expansion across different subtypes. (A) Bar plot showing the number of cells from the top 10 clonotypes. (B) Bar plot showing the number of cells from the top 10 clonotypes in MG and NoMG samples. (C) Clonal expansion observed in specific CD8+ T cell subtypes (CD8.Tem, CD8.Temra, and CD8.Trm), illustrated by the large number of cells belonging to several clonotypes.

**Figure S7.**
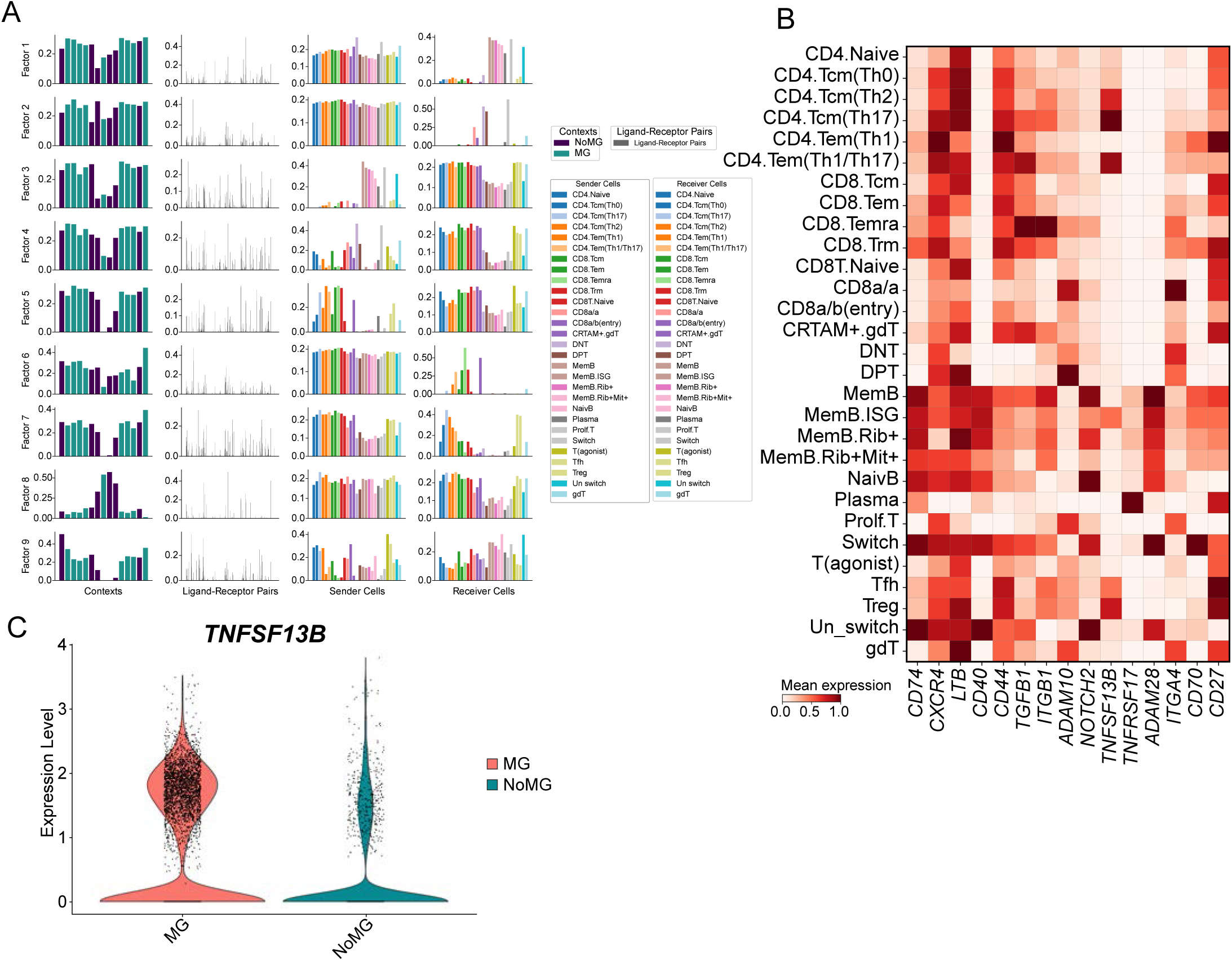
Altered T–B cell communication patterns in MG samples revealed by Tensor-Cell2cell analysis. (A) Bar plot illustrating the distribution of weight values for each factor derived from Tensor-Cell2cell analysis. (B) Heatmap showing the relative expression of key ligand–receptor gene pairs across cell subtypes. (C) Violin plot demonstrating the overexpression of *TNFSF13B* in myeloid cells from MG samples.

**Table S1:** Characteristics of the patients in this study.

| Patient # | Sample # | Specimen Type | Chemistry | Diagnosis | Antibodies | Age | Sex |
| --- | --- | --- | --- | --- | --- | --- | --- |
| PT1 | S1 | Thymoma w/o MG | Single Cell 3' v3 | Thymoma type B2. T1b. Masaoka stage 2a. No MG | / | 53 | F |
| PT2 | S2 | Thymus w MG | Single Cell 3' v3 | Benign thymic tissue with hyperplasia. MG. Ocular symptoms | High AchR antibodies elevated | 24 | F |
| PT3 | S3 | Thymus w/o MG | Single Cell 3' v3 | Thymic cyst and Thymic hyperplasia. No MG | / | 60 | M |
| PT4 | S4 | Thymoma w/o MG | Single Cell 3' v3 | No MG | / | 50 | F |
|  | S5 | Thymus w/o MG | Single Cell 3' v3 |  |  |  |  |
| PT5 | S6 | Thymus w MG | Single Cell 5' R2 plus V(D)J | Thymic hyperplasia, MG | AchR positive | 41 | F |
| PT6 | S7 | Thymus w MG | Single Cell 5' R2 plus V(D)J | Focal lymphoid hyperplasia. Ocular Myasthenia | AchR positive, | 42 | F |
| PT7 | S8 | Thymus w MG | Single Cell 5' R2 plus V(D)J | Thymic hyperplasia, MG | AchR negative, LRP4 positive | 37 | F |
| PT8 | S9 | Thymus w/o MG | Single Cell 5' R2 plus V(D)J | Thymoma B1 | / | 53 | F |
| PT9 | S10 | Thymus w MG | Single Cell 5' R2 plus V(D)J | Thymic MG; Thymoma B2B3; metastases | AchR positive | 39 | F |
|  | S11 | Drop Metastases | Single Cell 5' R2 plus V(D)J |  |  |  |  |
|  | S12 | Thymoma w MG | Single Cell 5' R2 plus V(D)J |  |  |  |  |
|  | S13 | Metastases continuity | Single Cell 5' R2 plus V(D)J |  |  |  |  |
| PT10 | S14 | Thymoma w/o MG | Single Cell 5' R2 plus V(D)J | Thymoma B1 | / | 78 | M |
| PT11 | S15 | Thymus w MG | Single Cell 5' R2 plus V(D)J | Ocular MG | AchR negative, LRP4 negative, musk negative | 27 | F |
| PT12 | S16 | Thymus w MG | Single Cell 5' R2 plus V(D)J | Type AB thymoma | AchR positive | 50 | M |
|  | S17 | Thymoma w MG | Single Cell 5' R2 plus V(D)J |  |  |  |  |
| PT13 | S18 | Thymoma w MG | Single Cell 5' R2 plus V(D)J | Thymoma AB. MG | AchR negative | 53 | M |
| PT14 | S19 | Thymic Carcinoma | Single Cell 5' R2 plus V(D)J | No MG | / | 71 | F |
|  | S20 | Thymoma (Metastatic) | Single Cell 5' R2 plus V(D)J |  |  |  |  |
| PT15 | S21 | Thymus w MG | Single Cell 5' R2 plus V(D)J | MG, normal thymus | AchR positive | 21 | F |
| PT16 | S22 | Thymus w/o MG | Single Cell 5' R2 plus V(D)J | MG negative | / | 71 | M |
|  | S23 | Thymic Carcinoma | Single Cell 5' R2 plus V(D)J |  |  |  |  |

